# Differential profiles of soluble and cellular toll like receptor 2 and 4 in chronic periodontitis

**DOI:** 10.1101/355347

**Authors:** H. AlQallaf, Y. Hamada, S. Blanchard, D. Shin, R.L. Gregory, M. Srinivasan

**Affiliations:** Department of Periodontics and Allied Dental Programs, School of Dentistry, Indiana University–Purdue University Indianapolis.; Department of Biomedical and Applied Sciences, School of Dentistry, Indiana University–Purdue University Indianapolis.; Department of Oral Pathology, Medicine and Radiology, School of Dentistry, Indiana University–Purdue University Indianapolis.

**Keywords:** Toll like receptor, periodontitis, biomarker, saliva

## Abstract

Chronic periodontitis is a common inflammatory disease initiated by a complex microbial biofilm and mediated by the host response causing destruction of the supporting tissues of the teeth. Host recognition of pathogens is mediated by toll-like receptors (TLRs) that bind conserved molecular patterns shared by groups of microorganisms. The oral epithelial cells respond to most periodontopathic bacteria via TLR-2 and TLR-4. Many studies have previously reported the presence of elevated numbers of viable exfoliated epithelial cells (SEC) in the saliva of patients with chronic periodontitis. In addition to the membrane-associated receptors, soluble forms of TLR-2 (sTLR-2) and TLR-4 (sTLR-4) have been identified and are thought to play a regulatory role by binding microbial ligands. sTLR-2 has been shown to arise from ectodomain shedding of the extracellular domain of the membrane receptor and sTLR-4 is thought to be an alternate spliced form. The objective of this study was to investigate the potential value of salivary sTLR-2/4 and the paired epithelial cell-associated TLR-2/4 mRNA as diagnostic markers for chronic periodontitis. Unstimulated whole saliva was collected after obtaining informed consent from 40 individuals in either periodontitis or gingivitis cohorts. The levels of sTLR-2/4 were measured by enzyme-linked immunosorbent assay (ELISA). SEC TLR-2/4 transcripts were quantitated by real time polymerase chain reaction. While levels of sTLR-2 exhibited an inverse correlation, sTLR-4 positively correlated with clinical parameters in the gingivitis cohort. Interestingly, both correlations were lost in the periodontitis cohort indicating a dysregulated host response. On the other hand, while sTLR-2 and the paired SEC associated TLR-2 mRNA exhibited a direct correlation (r^2^=0.62), that of sTLR4 and SEC TLR-4 mRNA exhibited an inverse correlation (r^2^=0.53) in the periodontitis cohort. Collectively, assessments of salivary sTLR2 and sTLR4 together with the respective transcripts in SECs could provide clinically relevant markers of disease progression from gingivitis to periodontitis.

## Introduction

Periodontitis is a microbial biofilm-induced, host-immune mediated, chronic inflammatory disease that leads to the destruction of the supporting apparatus of the teeth, if undiagnosed and untreated. Nearly 46% of US adults, ages 30 years and older, representing 64.7 million people, suffer from periodontitis, with 8.9% or 12.3 million having severe periodontitis (1). Changing composition of the dental plaque biofilm from healthy or symbiotic to a pathogenic flora promotes clinical progression of the disease from a periodontally healthy state to destructive periodontitis. Pathologically, the progression involves skewing of the host response from a predominantly protective innate immune response to an exaggerated response characterized by increased pro-inflammatory cytokines, eicosanoids, reactive oxygen species and matrix metalloproteinases that cause destruction of the tooth supporting tissues (2).

The main goal of periodontal therapy is to remove pathogenic biofilm and debride the affected tissues, to facilitate resolution of inflammation and healing of tissue attachment around teeth. Together with meticulous home care this should reduce bacterial accumulation (3). However, long-term periodontal health maintenance largely depends on regular and frequent periodontal maintenance. Current methods of diagnosing and monitoring periodontitis include measurement of probing depths, recessions, clinical attachment levels, bleeding on probing, presence of plaque, suppuration and radiographic bone loss. Yet, when used alone, these methods assess periodontitis only after the biologic onset of the disease process, and are unable to substantiate disease activity or future risk. Hence, over the years there has been an active search for identifying biomarkers as a supplemental diagnostic and risk assessment tool for managing periodontitis (4).

Host recognition of microorganisms is largely mediated by pattern recognition receptors (PRRs), such as toll like receptors (TLR) that recognize conserved molecular patterns, termed pathogen-associated molecular patterns (PAMPs) shared by groups of microorganisms. To date thirteen mammalian TLRs and many of their ligands have been identified. In addition to membrane-associated TLRs, soluble forms of TLRs (sTRLs) have been identified in serum, urine, tears and saliva (5–7). sTLRs are thought to act as negative regulators and inhibit TLR mediated signaling_pathways. Based on their ligand preferences, TLR-2 and TLR-4 respond to most periodontal pathogens by binding the peptidoglycan of the gram positive and the lipopolysaccharide of gram negative bacterial cell walls, respectively. Both, TLR-2 and TLR-4 work with co-receptor CD14 in binding periodontal pathogens. Altered expression profiles of CD14, TLR-2 and TLR-4 have been previously reported in periodontitis (8, 9). Additionally, periodontal pathogens have been shown to induce TLR-2/4 mediated signaling and up-regulate cytokine production in gingival epithelial cells, which ultimately leads to tissue destruction in periodontitis (10, 11).

Early investigations on biomarkers for periodontitis focused on the use of serum and gingival crevicular fluid in the assessment of inflammatory molecules. However, in the recent past considerable efforts have been made in identifying markers in saliva as it is an easily accessible biospecimen that is amenable for frequent and painless collection (12). Different classes of molecules such as cytokines, matrix metalloproteinases, and micro RNAs in saliva have been evaluated as biomarkers of periodontitis, however, the evidence of disease association has been inconsistent and difficult to repeat (13–16). Recently, levels of microbial ligands for TLR-2 and TLR-4 have been reported to be higher in saliva of periodontitis patients. Interestingly, many studies reported that sTLR-2 and sTLR-4 exhibited a decreasing trend in periodontitis saliva. (17)

The number of exfoliated salivary epithelial cells (SECs) have been shown to be higher in saliva of chronic periodontitis patients, supporting increased epithelial exfoliation secondary to inflammation (18). Interestingly, we observed that much like the reports in human gingival epithelial cells, stimulation with TLR-2 or TLR-4 specific ligands induced cytokine secretion with differential kinetics and up-regulated TLR-2 and TLR-4 mRNAs, respectively, in cultures of SECs from patients with periodontitis (19). This suggests that SECs could represent an excellent biological resource for investigating the host response to periodontal pathogens.

The purpose of this study was to detect potential correlations between soluble and epithelial cell associated expression of TLR-2 and TLR-4 in saliva of chronic periodontitis patients and their possible use as biomarkers to assess the status and progression of periodontal diseases.

## Materials and Methods

### Study population and clinical measurements

Prior to initiation of this study the Institutional Review Board at Indiana University Purdue University at Indianapolis (IUPUI) approved this protocol to access the saliva bank (IRB #1209009585). The study included 20 individuals that had gingivitis [minimal to no CAL, no radiographic evidence of bone loss] and 20 age matched individuals exhibiting clinical features of generalized moderate to severe chronic periodontitis [> 30% of sites having >4 mm of clinical attachment loss (CAL)] (20). Periodontal diagnoses were validated with radiographic evidence of bone loss with the help of recent full-mouth radiographs. A modified Schei ruler was used to measure bone loss, and an average measurement of the mesial and distal bone levels was obtained (21). The Ramfjord teeth were selected_for measurement, and if a tooth was not present, the tooth distal to it was utilized (22). Twelve radiographs were randomly selected and an inter/intra examiner measurement calibration was performed (HA and YH). An interclass correlation coefficient (ICC) > 0.9 was achieved prior to the actual measurements. Clinically, gingivitis controls should have minimal to no CAL, no overt clinical signs of gingival inflammation, and minimal bleeding on probing (BOP) scores, while the chronic periodontitis group should exhibit severe clinical inflammation, and high BOP scores. Patients with a diagnosis of aggressive periodontitis, any known systemic illness, history of routine use of antibiotics/anti-inflammatory therapy within the past 6 months, subjects with oral mucosal lesions and present or past history of smoking were excluded from both groups.

### Saliva collection and isolation of epithelial cells

Unstimulated whole saliva (UWS) was collected as follows. Following the measurement taking and subject recruitment visit, subjects refrained from eating or drinking for 1 h prior to saliva collection. They were seated with their head tilted towards one side. Subjects were asked to swallow and UWS was collected by a drooling method for 10 min into a 15 ml chilled centrifuge tube and transported on ice to the laboratory for processing. All UWS samples were centrifuged at 250 × g for 10 min at 4°C. Cellular sediment was obtained and reconstituted in a 1:10 volume of isotonic saline with an addition of two drops of Zap-O-globin to lyse blood corpuscles and centrifuged at 1,271.7 × g for 10 min at 4°C. After washing with saline the cell suspension was filtered through a membrane of 20-micron pore size. The membrane-trapped epithelial cell-enriched cell preparation was assessed by light microscopy for appropriate morphology, reconstituted in RPMI-1640 (Mediatech Inc., Minnesota, MN, USA), supplemented with 5% fetal bovine serum (Hyclone fetal bovine serum, Thermo Scientific, Logan, UT, USA) and 5% dimethylsulfoxide (Sigma-Aldrich, St Louis, MO, USA), and stored at -80°C until further analysis.

### ELISA for sTLR-2 and sTLR-4

Protein content of clarified saliva was determined by spectrophotometry using the Bradford method to increase the sensitivity of detection of low abundant proteins. Each UWS sample was depleted of amylase and immunoglobulins by incubating serially with anti-human amylase mAb (1:2500; Abcam, Cambridge, MA, USA) and protein G beads (Miltenyi Biotec Inc Auburn, CA) at 4°C. One microgram of protein from each treated UWS sample was assessed for the presence of sTLR-2/sTLR-4 by ELISA. sTLR-2 and sTLR-4 were detected in duplicate using anti-human TLR-2 mAb and anti-human TLR-4 mAb (R&D Systems), respectively. Bound antibodies were detected using HRP-conjugated anti-mouse immunoglobulin (Ig)G followed by TMB (3,3′,5,5′-tetramethylbenzidine) substrate (Pharmingen, San Diego, CA, USA). Purified recombinant human TLR-2Fc and TLR-4Fc (R&D Systems) was used to develop a standard curve. Absorbance at 450 nm was measured in a microplate reader (Biorad Laboratories, Hercules, CA, USA).

### Quantitative real-time PCR for epithelial TLR-2 and TLR-4

Total cellular RNA was isolated from SECs using a Qiagen RNA isolation kit (Invitrogen) and reverse-transcribed using an iScript cDNA synthesis kit (Biorad, Austin, TX, USA). An equal amount of cDNA was used to amplify TLR-2 and TLR-4 using SYBR green PCR master mix (SA Biosciences, Frederick, MA). Primers were designed using Primer Express software as follows: TLR-2 forward: 5’ACCTGTGTGACTCTCCATCC-3, reverse: 5’GCAGCATCATTGTTCTCTC-3; TLR-4 forward: 5’TTCCT CTCCTGCGTGAGAC-3’, reverse: 5’TTCATAGGGTTCAGGGACAG-3’ and small proline-rich protein 2a (SPRR2a) forward: 5AGTGCCAGCAGAAATATCCTCC-3, reverse: 5’GAACGAGGTGAGCCAAATATCC-3’ and glyceraldehyde dehydrogenase. TLR-2 and TLR-4 mRNA in SECs were amplified as positive controls. The PCR products were visualized and images acquired. ΔCt was obtained by subtracting the Ct for TLR-2 or TLR-4 from that of SPRR2a for each sample. The magnitude of change in the mRNA was expressed as 2^−ΔCt^ (19). Each measurement of a sample was conducted in duplicate.

### Statistical Methods

Statistical differences in the expression of TLRs between healthy and chronic periodontitis patients were determined using two-sample t-tests. The distribution of the expression levels was examined and a transformation of the data (e.g., natural logarithm) or nonparametric tests was used. Plots and Pearson correlation coefficients were used to test the association between soluble and cellular TLRs expression. A 5% significance level was used for all tests.

## Results

### Demographic and clinical features

The final study cohort included 20 gingivitis patients with an average age of 46.10 ± 11.55 years and 20 patients with chronic periodontitis with an average age of 51.25 ± 13.77 years (Table 1). The male to female ratio was 5:11 in gingivitis patients and 15:3 in periodontitis patients. Periodontal measurements, including plaque index, bleeding on probing and percentage of sites with pocket depth of >4 mm and mean clinical attachment levels of >4 mm were significantly higher in the periodontitis cohort. In the group with chronic periodontitis more than 50% of sites exhibited clinical attachment loss of >4 mm, and a 20.83 ± 6.41 percentage of bone loss, meaning that this cohort is a good representation of moderate chronic periodontitis.

**Table 1:** Clinical profile of gingivitis and chronic periodontitis subjects.

| Patient Profile | Gingivitis N = 20 (Mean ± SD ) | Periodontitis N =20 (Mean ± SD ) |
| --- | --- | --- |
| *Age (years)* | 46.10 ± 11.55 | 51.25 ±13.77 |
| *Gender ratio (M/F)* | (5:11) | (15:3) |
| *PD (mm) \** | 2.17 ± 0.321 | 3.55 ± 0.88 |
| *CAL (mm) \** | 2.22 ± 0.31 | 3.97 ± 1.01 |
| *BOP (%) \** | 4.77 ± 7.14 | 38.1 ± 24.15 |
| *PI (%) \** | 19.88 ± 30.7 | 63.28 ± 22.49 |
| *Sites of PD > 4 mm (%) \** | 2.11 ± 3.78 | 39.33 ± 22.47 |
| *Sites of CAL > 4mm (%) \** | 2.90 ± 3.92 | 51.98 ± 18.67 |
| *Average Bone loss (%) \** | 0.530 ± 0.81 | 20.83 ± 6.41 |

\* P ≤0.05

### sTLR-2 and sTLR-4 are lower in chronic periodontitis

PAMPs of TLR2 and TLR4 have been shown to be elevated in saliva of periodontitis patients. Since sTLRs are thought to function as decoy receptors for their respective PAMPs, we measured sTLR-2 and sTLR-4 in saliva. We observed that while both sTLR-2 and sTLR-4 showed a trend toward significance between both cohorts, only the reduction in sTLR-4 was significant (P ≤0.05; Fig. 1a & 2a).

**Figure 1:**
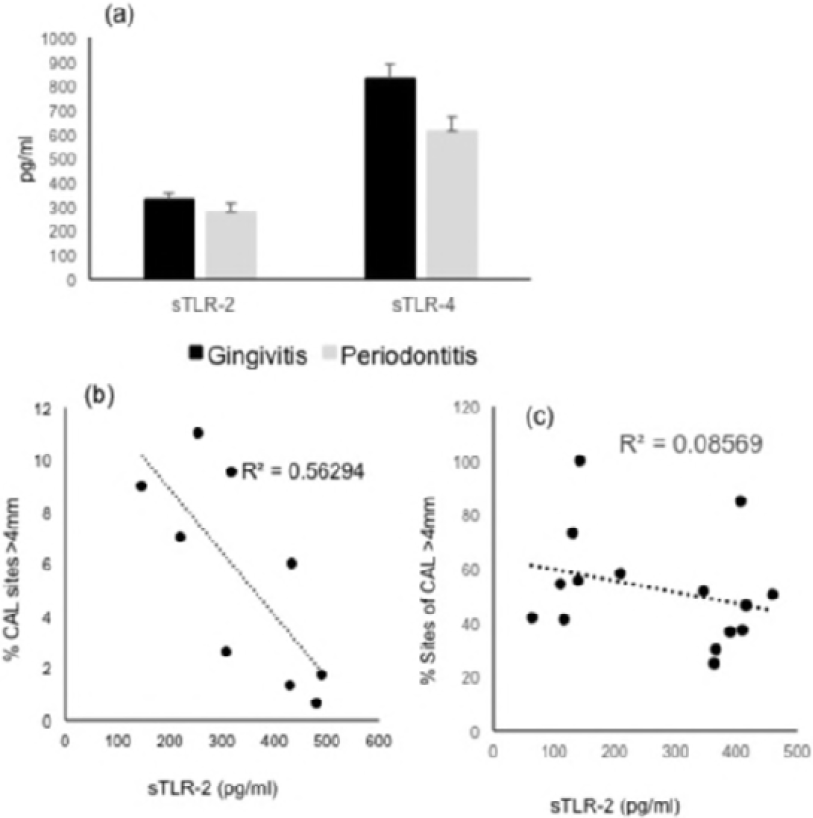
Relationship between sTLR-2 and clinical periodontitis. Unstimulated whole saliva (UWS) was collected from 20 patients with gingivitis and 20 patients with CP (defined as the presence of 30% of sites >4 mm CAL). (A) UWS sTLR-2 is lower in CP. Pearson correlation between the % CAL of >4mm and the paired sTLR-2 showed a moderate inverse correlation in the gingivitis cohort (B) and poor correlation in the CP cohort (C).

**Figure 2:**
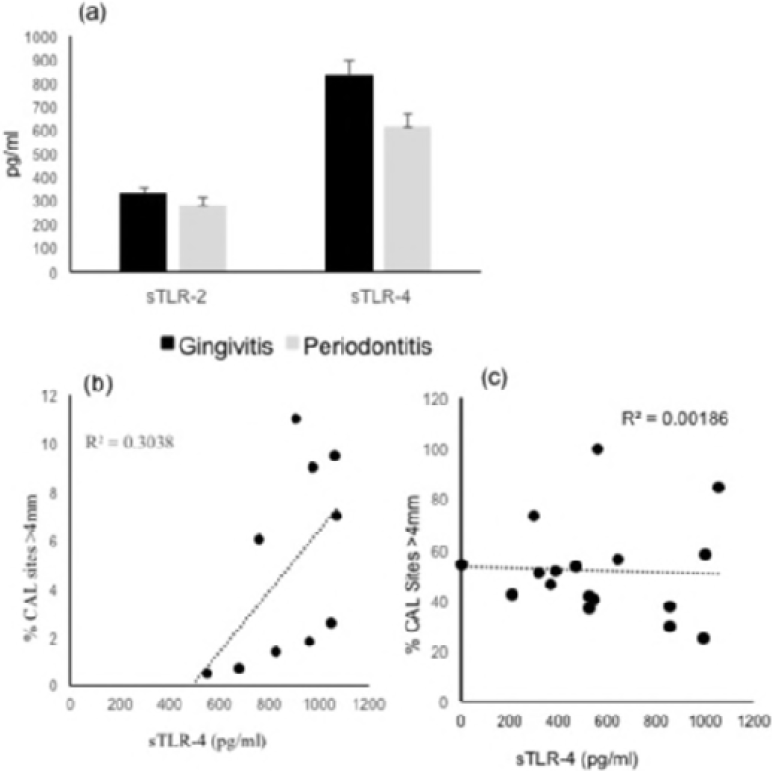
Relationship between sTLR-4 and clinical periodontitis. Unstimulated whole saliva (UWS) was collected from 20 patients with gingivitis and 20 patients with CP (defined as the presence of 30% of sites >4 mm CAL). UWS sTLR-4 is lower in CP. (A) Pearson correlation between the % CAL of >4mm and the paired sTLR-4 showed a moderate direct correlation in the gingivitis cohort (B) and poor correlation in the CP cohort (C).

### Correlation of sTLR-2 and sTLR-4 with clinical parameters

To evaluate whether sTLR-2 or sTLR-4 could be potential biomarkers for chronic periodontitis, we determined pairwise correlations of sTLR-2 and sTLR-4 with clinical parameters. Interestingly, an inverse correlation was observed between sTLR-2 and paired clinical parameters of % CAL (R^2^=0.56) or % bleeding index (R^2^=0.63) in the gingivitis cohort (Fig. 1b). In contrast, we observed a direct correlation between sTLR-4 and the clinical parameters in the gingivitis cohort with higher sTLR-4 correlating with a higher % CAL and % bleeding index (R^2^=0.3; Fig 2b). However, neither sTLR-2 nor sTLR-4 in saliva correlated with the paired %CAL or % bleeding index in the periodontitis cohort (Fig 1c & 2c). Taken together, these results suggest that sTLR-2 and sTLR-4 could potentially represent markers of gingivitis prior to bone loss.

### Epithelial cell expression of TLR-2 and TLR-4 (Fig. 3a & 4a)

The TLR-4 transcript in SECs is up-regulated in chronic periodontitis. Previously, we observed that the relative expression of TLR-4 mRNA from epithelial cells isolated from periodontitis saliva is higher than that isolated from healthy patient saliva. (19) In this study, we observed similar higher TLR-4 mRNA but equivalent levels of TLR-2 mRNA in SECs in saliva from chronic periodontitis patients as compared to that from gingivitis patients.

**Figure 3:**
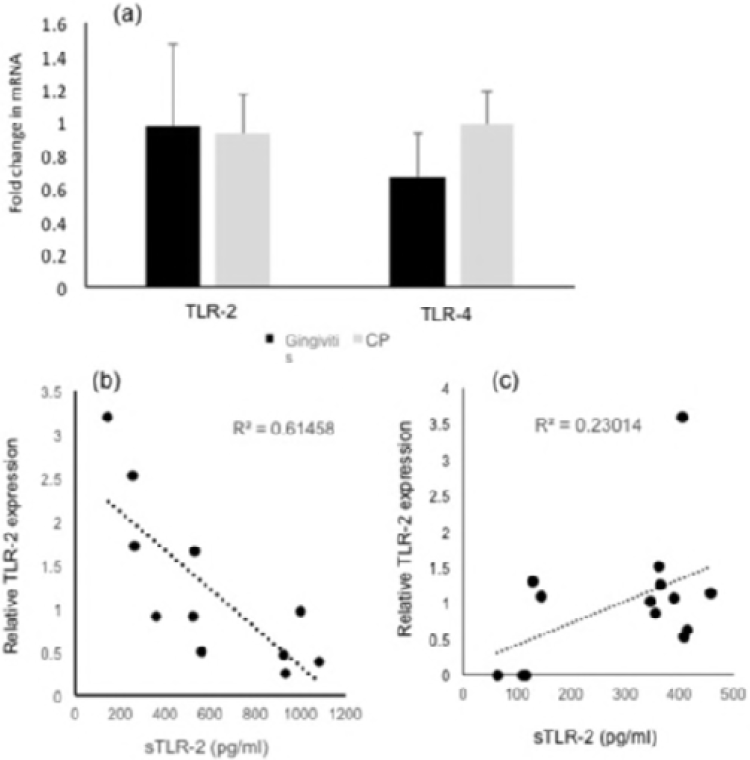
Relationship between sTLR-2 and SECs associated TLR-2 transcript. Unstimulated whole saliva (UWS) was collected from 20 patients with gingivitis and 20 patients with CP (defined as the presence of 30% of sites >4 mm CAL). UWS were processed to separate clarified and epithelial cell rich fraction as described in materials and methods. The concentration of sTLR-2 was measured by ELISA. The SEC associated TLR-2 mRNA was determined by quantitative PCR. (A) Relative quantities of TLR-2/TLR-4 mRNA with respect to SPRR2a mRNA were determined using the 2^−ΔCt^ method. (B) Shows a moderate inverse correlation between sTLR-2 and SEC TLR-2 mRNA in the gingivitis cohort and (C) shows a mild direct correlation between the sTLR-2 and SEC TLR-2 mRNA in the periodontitis cohort.

**Figure 4:**
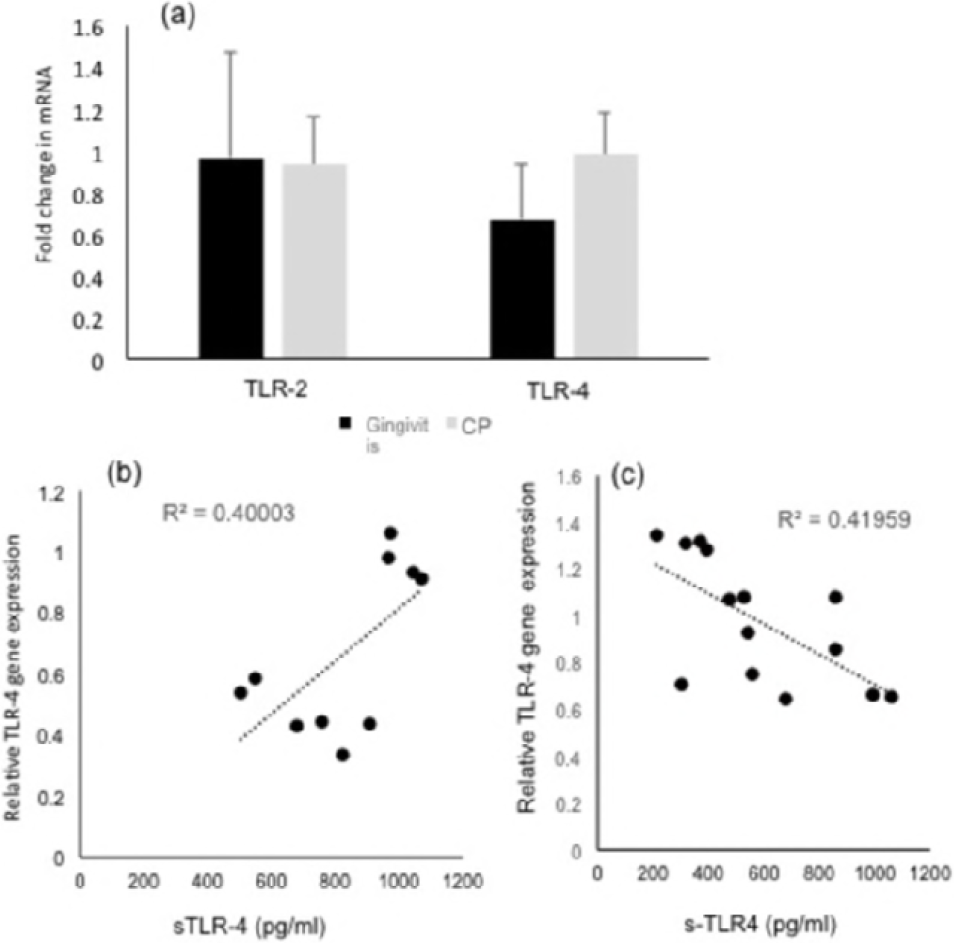
Relationship between sTLR-4 and SECs associated TLR-4 transcript. Unstimulated whole saliva (UWS) was collected from 20 gingivitis individuals and 20 patients with CP (defined as the presence of 30% of sites >4 mm CAL). UWS were processed to separate clarified and epithelial cell rich fraction as described in materials and methods. The concentration of sTLR-4 was measured by ELISA. The SEC associated TLR-4 mRNA was determined by quantitative PCR. (A) Relative quantity of TLR-2/TLR-4 mRNAs with respect to SPRR2a mRNA was determined using the 2^−ΔCt^ method. (B) Shows a moderate positive correlation between sTLR-4 and SEC TLR4 mRNA in the gingivitis cohort and (C) shows a moderate inverse correlation between the sTLR-4 and SEC TLR-4 mRNA in the periodontitis cohort.

### Correlation of sTLR-2/sTLR-4 with SEC associated TLR-2/TLR-4

Since sTLRs are thought to arise from extracellular shedding of membrane associated TLR, we next investigated whether sTLR-2 and sTLR-4 levels correlate with SEC associated TLR-2 or TLR-4. Evaluation of the relationship between sTLR-2 with paired SEC TLR-2 mRNA in the gingivitis cohort suggested a moderate inverse correlation (Fig 3b). However, there was a mild direct correlation between sTLR-2 and paired SEC TLR-2 mRNA in periodontitis saliva (Fig 3c). In gingivitis saliva, lower concentrations of sTLR-4 correlated with lower TLR-4 mRNA levels in paired SEC (Fig 4b). As opposed to this, in periodontitis saliva, sTLR-4 concentrations exhibited a moderate inverse correlation with paired SEC associated TLR-4 mRNA levels (Fig 4c). Taken together these observations, the negative correlation between sTLR-2 levels and paired SEC TLR-2 mRNA in health, provides additional evidence of the origin of sTLR-2 by shedding of the ectodomain of the membrane bound TLR-2.

## Discussion

TLR function as important signal transducers that mediate innate immune and inflammatory responses to pathogens through pattern recognition of virulence molecules. Almost all human cells express a unique ratio of TLRs that allows them to respond to a plethora of microorganisms (23). Optimal TLR activation and regulated response is critical to maintaining a healthy host-microbe homeostasis. Dysregulated activation of TLRs is directly involved in the pathogenesis of chronic microbial diseases including periodontitis (24). Multiple intracellular and extracellular regulatory mechanisms have been shown to balance TLR dependent responses. sTLRs acting as decoy receptors represent one of the mechanisms of negative regulation. sTLRs are thought to bind to and directly attenuate PAMPs in the extracellular space preceding their engagement with specific PRRs. To date, four extracellular sTLRs have been identified in humans, including sTLR-1, sTLR-2, sTLR-4, and sTLR-6. Of these four extracellular sTLRs, sTLR-2 and sTLR-4 have been detected in most body fluids including saliva, serum, urine and breast milk (5–7).

sTLR-2 has been shown to be elevated in many chronic inflammatory conditions, such as inflammatory bowel diseases, HIV infection, and various cardiovascular conditions (23, 24). It originates by proteolytic cleavage of the TLR-2 transmembrane protein through a process of ectodomain shedding mediated by metalloproteinases (25). Since metalloproteinases are up-regulated in chronic periodontitis, production of sTLR-2 would serve to diminish detrimental inflammation (26). It has been suggested that during innate immune responses, ectodomain shedding is a strategy that permits downregulation of responses triggered by pathogens or stressors (23, 24). Furthermore, studies in mice suggest that sTLR-2 significantly reduces bacteria-associated inflammation without impairing microbial clearance (27). Pertinently, it is interesting that our data demonstrates a negative correlation between SEC TLR-2 mRNA and sTLR-2 in the gingivitis cohort and not in the periodontitis cohort. This suggests that sTLR2 and the SEC associated TLR-2 transcript may represent markers for periodontal health. Previously, in monocytes microbial stimulation mediated negative correlation between membrane-bound TLR-2 and sTLR-2 in culture supernatant.

Several studies have reported the expression and activation of TLR-4 in chronic periodontitis. Banu et al. reported an increase in sTLR-4 in saliva as opposed to our observation of lowered sTLR4 (28). The discrepancy can be attributed to differences in the nature of the sample (stimulated versus unstimulated saliva), disease status at the time of sample collection and the use of a different assay kit with potential differences in the sensitivity and specificity of the antibodies in the kit. Higher plasma TLR-4 has been reported in periodontitis. Although naturally sTLR-4 has been suggested to arise as alternate splice variant, it is also possible that passive diffusion could contribute to the overall concentration of sTLR-4 in saliva (28, 29). Lappin et al. reported elevated levels of TLR-2 and TLR-4 stimulants in periodontitis saliva (30). It has been suggested that sTLR-4 bound in a complex with MD2, a protein associated with TLR-4 on the host cell surface, may block the interaction between membrane-bound TLR-4 and its ligands and thereby inhibit TLR-4 signaling (24, 31). The lower sTLR-4 and the level of higher SECs TLR-4 mRNA in periodontitis saliva in our cohort could potentially reflect active sequestration of microbial ligands by sTLR-4 and compromised ability to block epithelial TLR-4. Furthermore, the direct correlation between higher sTLR-4 and greater SECs TLR4 mRNA in gingivitis saliva perhaps support the sentinel functions of TLR4.

It is well recognized that in chronic periodontitis the host response associated with bacterial invasion directs the disease progression from gingivitis (inflammation) to periodontitis (infection). While sTLR-2 has been suggested as a maker of inflammation in many conditions including colitis and arthritis, sTLR-4 has been suggested as an infection marker, although it may not correlate with microbial clearance (24, 27).

In this context, our data of relatively higher sTLR-2 in gingivitis and significantly lower sTLR-4 in periodontitis saliva suggest that assessment of both could provide a clinically viable diagnostic marker, a method to track disease progression and even response to therapy. Utilizing saliva to identify and measure specific phenotypes and host-derived mediators will allow highly individualized diagnosis, prognosis and treatments for periodontal diseases. This personalized medicine approach will strengthen the power of the clinical oral examination and medical history assessments, providing patients with evidence-based, targeted risk care.

